# Microbial ecology of coastal northern Gulf of Mexico waters

**DOI:** 10.1101/2023.11.17.567634

**Authors:** Michael W. Henson, J. Cameron Thrash

## Abstract

Estuarine and coastal ecosystems are of high economic and ecological importance, owing to their diverse communities and the disproportionate role they play in carbon cycling, particularly in carbon sequestration. Organisms inhabiting these environments must overcome strong natural fluctuations in salinity, nutrients, and turbidity as well as numerous climate change-induced disturbances such as land loss, sea level rise, and, in some locations, increasingly severe tropical cyclones that threaten to disrupt future ecosystem health. The northern Gulf of Mexico (nGoM) along the Louisiana coast contains dozens of estuaries, including the Mississippi-Atchafalaya River outflow, which dramatically influence the region due to their vast upstream watershed. Nevertheless, the microbiology of these estuaries and surrounding coastal environments have received little attention. To improve our understanding of microbial ecology in the understudied coastal nGoM, we conducted a 16S rRNA gene amplicon survey at eight sites and multiple timepoints along the Louisiana coast and one inland swamp spanning freshwater to high brackish salinities, totaling 47 duplicated Sterivex (0.2-2.7 µm) and prefilter (> 2.7 µm) samples. We cataloged over 13,000 ASVs from common freshwater and marine clades such as SAR11 (Alphaproteobacteria), *Synechococcus* (Cyanobacteria), and acI and *Candidatus* Actinomarina (Actinobacteria). We observed preferences for freshwater or marine habitats in many organisms and also characterized a group of taxa with specialized distributions across brackish water sites, supporting the hypothesis of an endogenous brackish-water community. Additionally, we observed brackish-water preferences for several aquatic clades typically considered marine or freshwater taxa, such as SAR11 subclade II, SAR324, and the acI Actinobacteria. The data presented here expand the geographic coverage of microbial ecology in estuarine communities, help delineate the autochthonous and allochthonous members of these environments, and provide critical aquatic microbiological baseline data for coastal and estuarine sites in the nGoM.

## Introduction

Estuarine environments are highly diverse, interconnected ecosystems that are exposed to strong natural fluctuations in salinity and nutrient availability (Crump et al. 1998, 1999; Campbell and Kirchman 2012). Salinity is known to be a major factor influencing microbial community structure and metabolic capacity (Lozupone and Knight 2007; Logares et al. 2009; Walsh et al. 2013; Dupont et al. 2014). Regulatory, functional, and metabolic differences between closely related microbial taxa underpin preferences for different salinity regimes (Logares et al. 2010; Salcher et al. 2011; Dupont et al. 2014; Henson et al. 2018b; Cabello-Yeves et al. 2022; Jurdzinski et al. 2023). These important differences can prevent microorganisms from circumventing the marine-freshwater threshold, resulting in infrequent transitions between these environments within taxonomic clades (Logares et al. 2009, 2010; Dupont et al. 2014; Henson et al. 2018b; Cabello-Yeves et al. 2022; Lanclos et al. 2023) and lower species diversity in intermediate salinities (Olli et al. 2019), as first hypothesized in the Remane curve (Remane 1934).

Under future climate scenarios, estuaries are expected to be influenced by elevated land loss, as well as changes to salinity, pH, temperature, and other factors (Nicholls et al. 2007; Elliott et al. 2019; Reed et al. 2020; Jurdzinski et al. 2023), which could fundamentally alter the microbial community composition and ultimately the processing of carbon and other nutrients (Dupont et al. 2014). Previous work has sought to investigate if brackish environments host autochthonous microbial communities uniquely adapted to these fluctuating ecosystems. Indeed, brackish bacterial communities are genomically and taxonomically differentiated from their freshwater and marine counterparts (Campbell and Kirchman 2012; Larsson et al. 2014; Hugerth et al. 2015; Celepli et al. 2017; Xia et al. 2017; Paver et al. 2018; Vidanage et al. 2020; Rasmussen et al. 2020; Cabello-Yeves et al. 2022; Mohapatra et al. 2023; Lanclos et al. 2023; Jurdzinski et al. 2023). For instance, taxa within the dominant clades *Synechococcus* and SAR11, specifically subcluster 5.2 and subclade III, respectively, have unique brackish ecotypes and corresponding genomic capacities such as pigment type, nitrogen utilization, and osmotic regulation that may have facilitated their transition into these dynamic environments (Larsson et al. 2014; Hugerth et al. 2015; Celepli et al. 2017; Lanclos et al. 2023; Jurdzinski et al. 2023). Moreover, because of the strong selection factors influencing marine-freshwater transitions (Logares et al. 2009), genomic plasticity may be critical to these brackish-adapted taxa to maintain abundances and large biogeographic distribution at intermediary salinities (Jurdzinski et al. 2023). Therefore, estuarine microorganisms can be phylogenetically distinct and possess unique genomic capacities to adapt to natural fluctuations in environmental conditions (Hugerth et al. 2015; Jurdzinski et al. 2023). However, the number of brackish ecosystems studied has been limited (Jurdzinski et al. 2023), necessitating the collection of more widespread baseline microbial community data to understand the impacts of future climate scenarios on the biogeographic distribution and functions of resident microorganisms.

The northern Gulf of Mexico (nGoM) provides an excellent coastal/estuarine study system because of its economic and ecological value (Rabalais et al. 1996; Adams et al. 2004; Lindstedt 2005; Vörösmarty et al. 2009). Influenced by two major rivers -- the Mississippi and Atchafalaya Rivers-- and a vast network of interconnected wetlands, the nGoM coastline is subject to continuous fluctuations in environmental conditions such as salinity, nutrients, and turbidity (Bianchi et al. 1999; Rabalais et al. 2002; King et al. 2012). Over the past century, the nGoM has lost an estimated 5000 km^2^ of land due to erosion (Nittrouer et al. 2012). Moreover, ecosystem models predicting future climate scenarios suggest continued loss of freshwater wetlands due to saltwater intrusion (Nittrouer et al. 2012; Reed et al. 2020) which will impact ecosystem functions across the coastal area, including biogeochemical cycling by microorganisms. However, previous microbiological research in this region has mostly focused on the communities associated with oil spills and eutrophication, and there is much less data on the microbial ecology separate from these stressors (Olapade 2010; King et al. 2012; Mason et al. 2012; Tolar et al. 2013; Gillies et al. 2015; Thrash et al. 2017, 2018; Noirungsee et al. 2020) limiting our ability to infer how the estuarine bacterioplankton in the nGoM naturally fluctuate over time in these diverse habitats.

To characterize the baseline microbial communities that inhabit the nGoM along the Louisiana coast and contribute more broadly to understanding estuarine and brackish water communities globally, we collected surface water at nine coastal, estuarine, and freshwater sites over multiple years and seasons. We quantified the particle-associated (> 2.7 µm) and planktonic microbial (0.2 - 2.7 µm) communities as well as the associated water chemistry. Six sites were sampled once a year for three years as part of our nGoM cultivation campaign to compare our isolates to the natural communities, but a comprehensive ecological analysis of these samples was not previously completed. (Henson et al. 2016, 2020). Two other sites were sampled three times over five months (Tables 1 and S1), and one site, an inland swamp, was sampled only once (Tables 1 and S1). Our results showcase the diverse communities of microbes inhabiting these environments that are adapted to different salinity regimes. While numerous clades show typical marine and freshwater preferences, other taxa displayed a euryhaline distribution that supports the hypothesis of a core group of autochthonous taxa uniquely adapted to brackish environments (Dupont et al. 2014; Hugerth et al. 2015; Cabello-Yeves et al. 2022; Lanclos et al. 2023; Jurdzinski et al. 2023). This study is a comprehensive description of the diverse microbial communities from nGoM estuarine and coastal systems and expands our biogeographic understanding of important and poorly characterized clades from these environments.

## Results

We sampled nine sites (47 samples size-fractionated at 0.22-2.7 µm [24 samples] and > 2.7 µm [23 samples]) from across southern Louisiana’s coastal, estuarine, and swamp environments over three years from 2014-2017 (**Fig. 1A**; **Table 1**). We found positive, significant correlations between nitrate (NO_3_^−^) and phosphate (PO_4_^2-^) (R=0.649,p=001) and nitrite (NO ^−^) and ammonium (NH_4_ ^+^) (R=0.618, p=0.003) (**Table S1:** Nut_Cor), while temperature and dissolved oxygen (DO) (R=-0.696, p<0.005) and salinity and silicic acid [Si(OH)_4_] were significantly, negatively correlated (R=-0.563, p<0.001) (**Table S1:** Nut_Cor). Sites ranged from fresh (< 0.5 salinity) to high brackish, i.e., polyhaline waters (max salinity=26.01 at JLB) (**Fig. 1A**). Principal component analysis of environmental conditions at the nine sites showed a distinct horizontal and vertical separation (**Fig. 1B**). Salinity and NO_2_ ^−^ were the most important values separating the sites. Sites above the 0 horizontal line typically had higher salinities (> 12) and higher NO_2_ ^−^ and NH_4_ ^+^ concentrations, while sites below the zero horizontal line were more indicative of high Si(OH)_4_ (**Fig. 1B**). Sites to the left of the vertical 0 line sites were indicative of lower pH and DO, while sites to the right typically had higher nitrate and phosphate concentrations (**Fig. 1B**).

**Figure 1.**
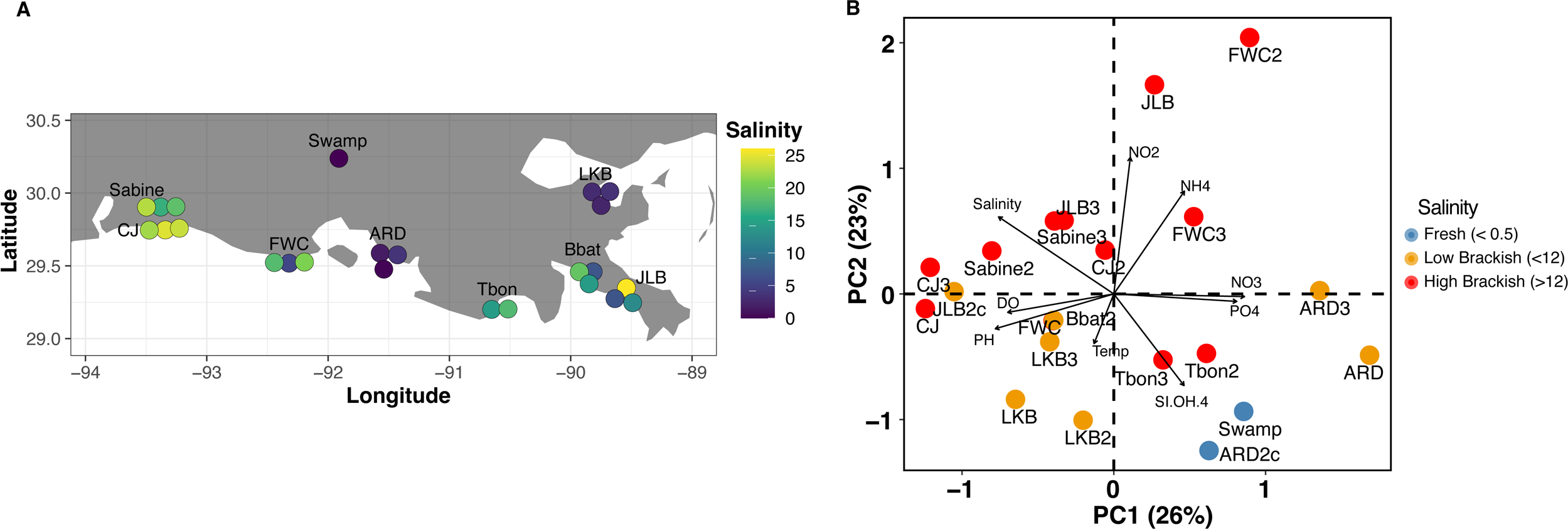
A) Locations of the nine sampling sites along the coast of the northern Gulf of Mexico. The dot color corresponds to the salinity at each site. The map was made with the R package ggplot2 using the command map_data. B) Two-dimensional principal coordinates analyses plot of normalized water characteristic variables measured at each site. Eigenvectors are scaled to strength. The percent variation each principal component explains is indicated in parentheses adjacent to the component axis.

**Table 1.** Site environmental measurements and location.

| Site | Temperature Range (°C) | Salinity Range | 0.2-2.7 $\mu\text{m}$ | > 2.7 $\mu\text{m}$ | Sampling Dates | Location | NOTES |
| --- | --- | --- | --- | --- | --- | --- | --- |
| *ARD* | 8.76-24.7 | 0.18-3.72 | published* | unpublished | 7/2015,<br>6/2016,<br>12/2016 | 29.575,<br>-91.538 | VERMILLION BAY,<br>ATCHAFALAYA RIVER DELTA<br>(BURNS POINT, LA) |
| *BBAT* | 20.61-30.42 | 6.92-19.09 | unpublished | unpublished | 7/2015,<br>9/2015,<br>11/2015 | 29.458,<br>-89.815 | BAY BATISTE<br>(PORT SULPHUR, LA) |
| *CJ* | 25.54-31.47 | 22.16-24.63 | published* | unpublished | 9/2014,<br>9/2015,<br>9/2016 | 29.760,<br>-93.340 | CALCASIEU JETTIES<br>(CAMERON, LA) |
| *FWC* | 19.91-22.9 | 5.39-20.9 | published* | unpublished | 3/2015,<br>4/2016,<br>11/2016 | 29.530,<br>-92.326 | FRESHWATER CITY, LA |
| *JLB* | 7.66-27.08 | 6.89-26.01 | published* | unpublished | 1/2015,<br>5/2015,<br>1/2017 | 29.348,<br>-89.538 | BAY POMME D'OR (BURAS, LA) |
| *LKB* | 19.29-30.15 | 2.39-3.55 | published* | unpublished | 6/2015,<br>7/2016,<br>2/2017 | 30.003,<br>-89.826 | LAKE BORNE, SHELL BEACH, LA |
| *Sabine* | 25.81-30.42 | 16.38-22.96 | unpublished | unpublished | 7/2015,<br>9/2015,<br>10/2015 | 29.920,<br>-93.381 | SABINE NATIONAL WILDLIFE REFUGE<br>(HUCKBERRY, LA) |
| *Swamp* | 17.12 | 0 | unpublished | unpublished | 11/2014 | 30.221,<br>-91.906 | LAKE MARTIN (BEAUX BRIDGE, LA) |
| *TBON* | 29.64-31.63 | 14.2-17.7 | published* | unpublished | 8/2015,<br>7/2016 | 29.207,<br>-90.647 | TERREBONNE BAY<br>(COCODRIE, LA) |

\* (Henson et al. 2016, 2018b, 2020)

Non-metric multidimensional analysis (NMDS) of the 47 communities showed three distinct groupings based on salinity: fresh (< 0.5 salinity), low brackish (0.5-12), and high brackish (< 12 salinity) (**Fig. 2**; NMDS stress 0.135; ANOSIM R=0.703, p=0.001). Filter fraction difference (0.2-2.7 vs. >2.7 µm) was significant but had low explanatory power (ANOSIM R=0.244, p=0.001). Moreover, salinity (R^2^=0.807, p=0.001) and silicic acid [Si(OH)_4_] (R^2^=0.563, p=0.001) were the two strongest environmental variables correlated to the NMDS ordination of the combined fraction analysis (**Fig. 2**; **Table S1**: envfit).

**Figure 2.**
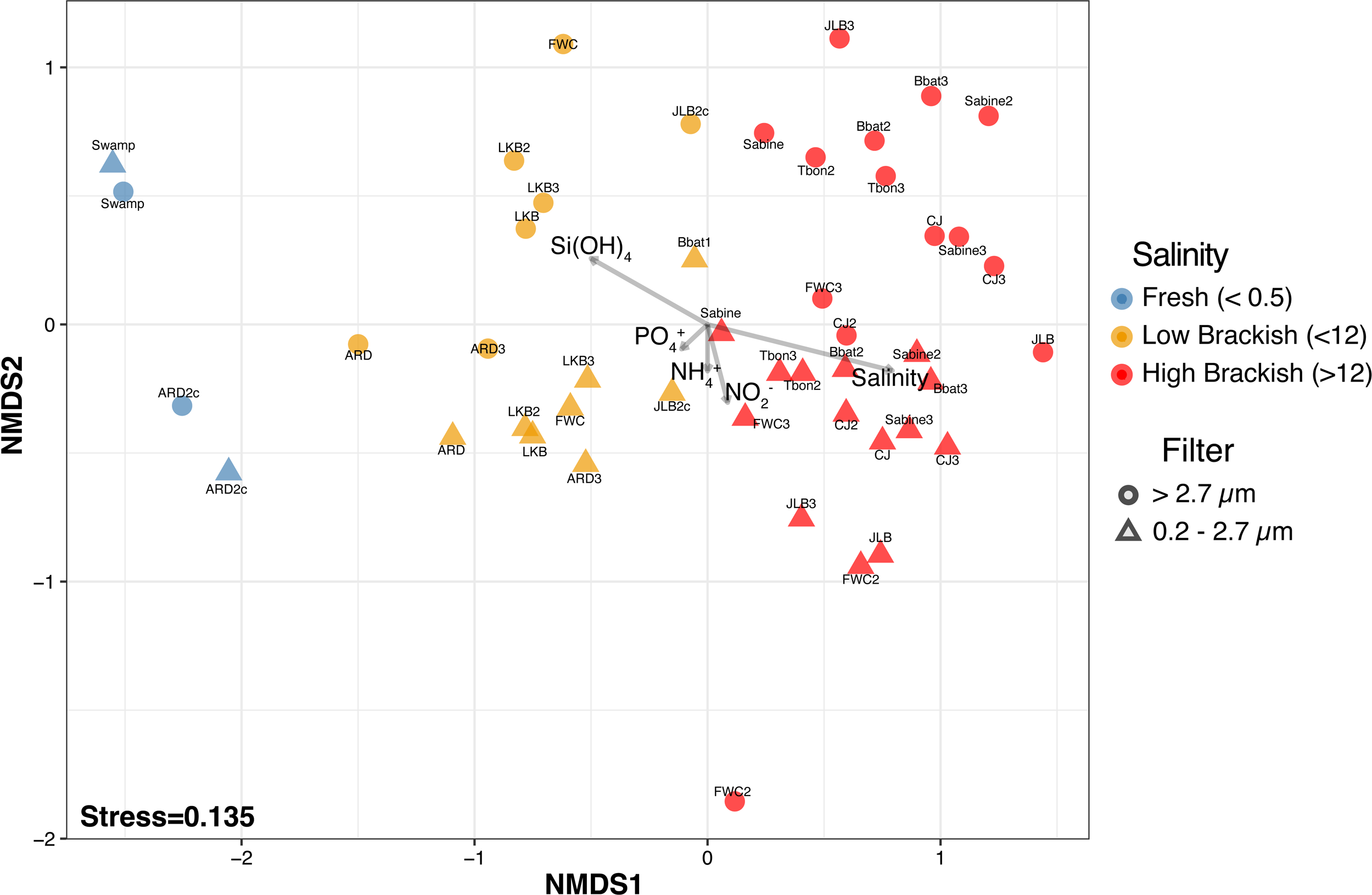
Non-metric multidimensional scaling ordination of the microbial communities of the nine sites. The dot color indicates the broad salinity classification of fresh (Blue, < 0.5 salinity), low brackish (orange, < 12 salinity), and high brackish (orange, > 12 salinity) at the sampling site. The dot shape indicates the community size fraction: triangle (0.2 - 2.7 µm) and circle (> 2.7 µm). Significant environmental variables (p < 0.05) are plotted as vectors. Arrow lengths have been adjusted based on their strength of correlation (R^2^).

### 0.2-2.7 µm associated communities

The 0.2-2.7 µm fraction communities examined separately from the > 2.7 µm fraction were strongly differentiated by salinity (R^2^=0.842, p=0.001), silicic acid (R^2^=0.475, p=0.016), and temperature (R^2^=0.386, p=0.005) (**Table S1**: envfit). The majority of reads in the 0.2-2.7 µm fraction classified to the clades SAR11 (Alphaproteobacteria), OM43 (Betaproteobacteria), *Candidatus* Actinomarina and acI (Actinobacteria), and *Cyanobium* (*Synechococcus* subcluster 5.2, Cyanobacteria) (**Fig. 3A**, **Table S1**: 0.2-2.7 µm RA). At higher salinities (> 12), typical marine clades such as SAR11 subclade I (Alphaproteobacteria) and *Candidatus* Actinomarina (OM1 clade, Actinobacteria) were more abundant than in samples of lower salinity (**Fig. 4A**). In contrast, typical freshwater clades such as SAR11 subclade IIIb/LD12 (*Candidatus* Fonsibacter sp., Alphaproteobacteria), and acI and acIV (Actinobacteria) were more abundant in lower salinities (**Figs. 4A**, **S1**; **Table S1**: 0.2-2.7 µm RA). Other ASVs with differentiated abundances were from the SAR86 (Gammaproteobacteria) and SAR324 clades, which both had higher relative abundance in higher salinity, while ASV54 unknown *Holophagaceae* (Acidobacteria) and ASV13 *Candidatus* Aqualuna (Actinobacteria) showed a significant preference for freshwater habitats (**Fig. 4A**, **Table S1**: 0.2-2.7 µm RA).

**Figure 3.**
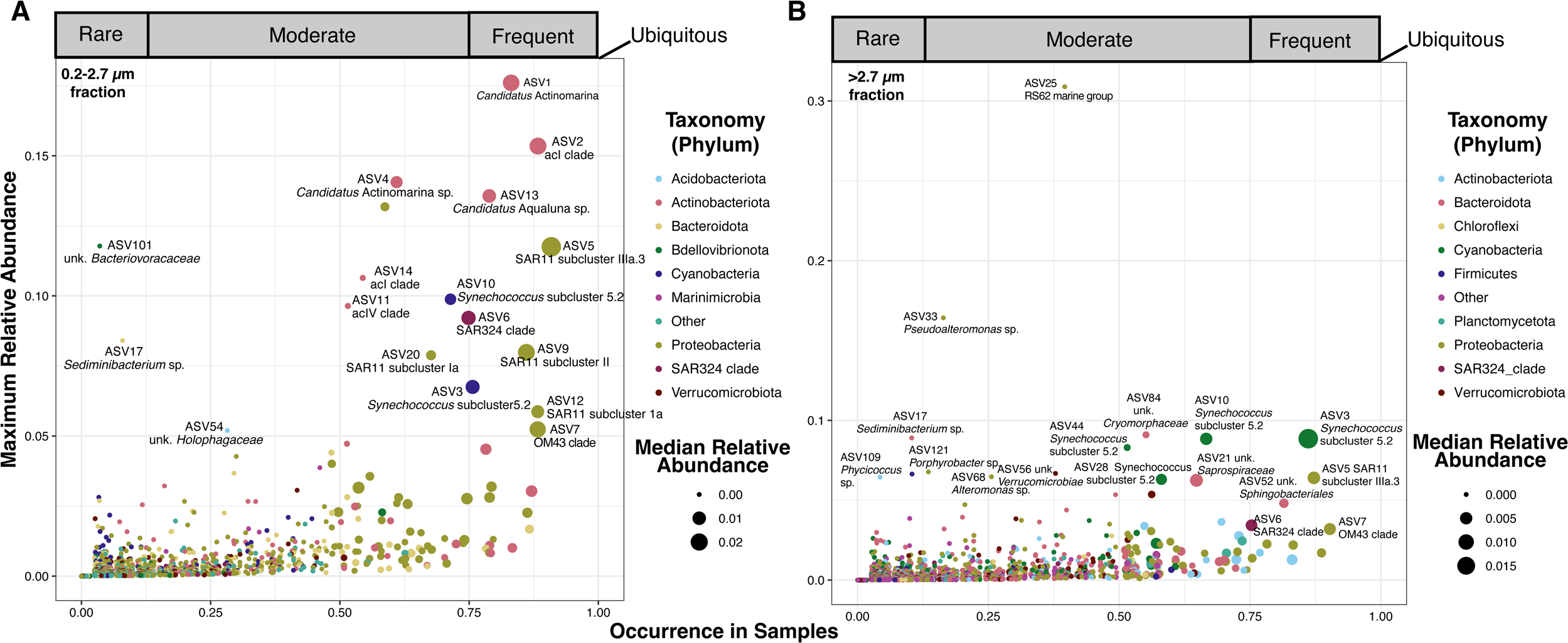
Maximum relative abundance of each ASV in the 0.2-2.7 µm (A) and > 2.7 µm (B) fraction according to its percent occurrence (>0%) across samples and maximum relative abundance. ASVs are color-coded by phylum and the size of the dot corresponds to the median relative abundance for each ASV.

**Figure 4.**
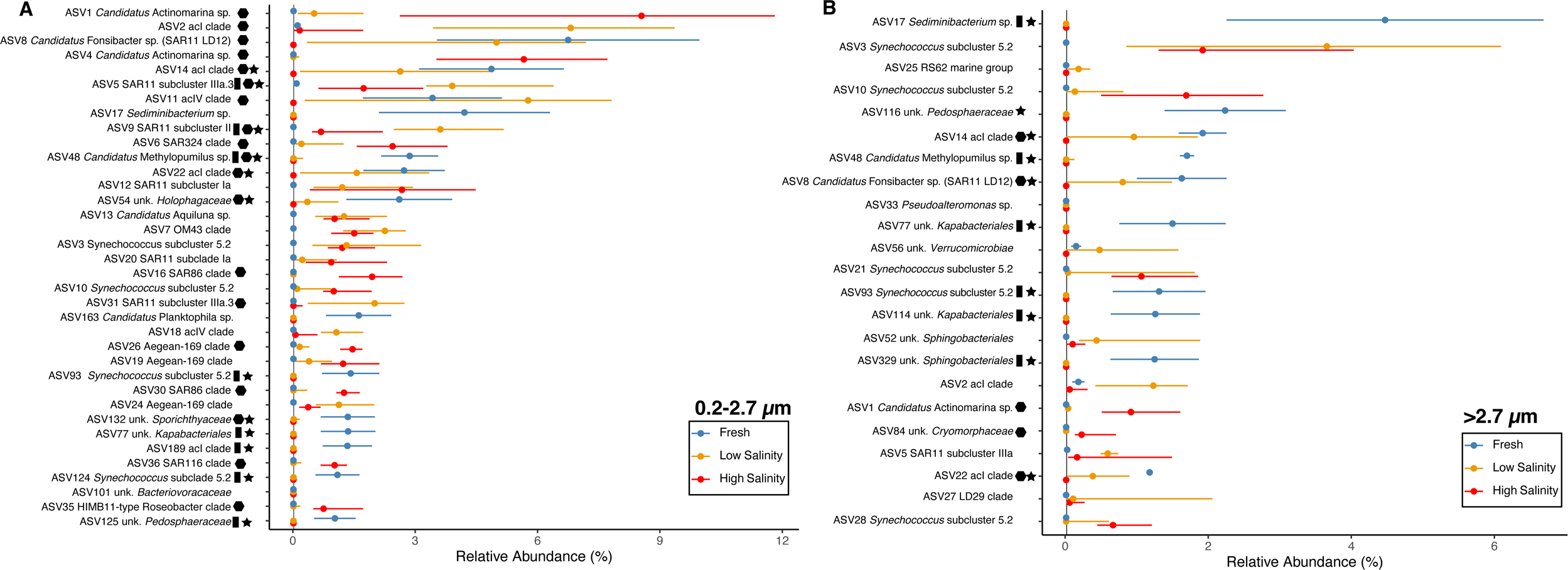
ASVs (> 1% RA) with significant differential abundance in fresh (Blue, < 0.5 salinity), low salinity (orange, < 12 salinity), or high salinity (orange, > 12 salinity) in the 0.2-2.7 µm (A) and > 2.7 µm (B) fractions according to the non-parametric, one way ANOVA on ranks (Krustal Wallis Test). Shape (Polygon=High Salinity, Low Salinity; Rectangle=Low Salinity, Fresh; Star=High Salinity, Fresh) indicate significant (p < 0.05) pairwise statistical differences between salinity conditions controlling for false discovery rate (Wilcoxon Test) (Table S1). Points represent median values, and lines represent the interquartile range. The vertical line indicates the limit of detection.

Taxa within the SAR11 and *Synechoccous* clades had diverse salinity habitat preferences (**Figs. 4A**, **S1**). ASVs from SAR11 subclade I increased in abundance with increasing salinity, while we observed subclade IIIa ASVs distributed more broadly across all brackish environments (**Figs. 4A, 5A, S1**; **Table S1**: SAR11 RA; **Table S1**: Pairwise). For instance, we observed ASV5 (SAR11 subclade IIIa.3) at relative abundances > 1% at sites ranging from low (> 0.5) to the maximum salinity sampled (26.01), overlapping with subclade I, while ASV31 (SAR11 subclade IIIa.3) occurred predominantly in lower salinity sites. We observed SAR11 subclade II-associated ASVs (e.g., ASV9 and ASV119) at high abundances across a breadth of salinities, complementing and expanding on previous descriptions of its distribution in aquatic habitats (Carlson et al. 2009; Vergin et al. 2013; Herlemann et al. 2014; Haro-Moreno et al. 2019; Lanclos et al. 2023) (**Figs. 5A, S1**; **Table S1**: SAR11 RA). *Synechococcus* ASVs were predominately from subcluster 5.2, with only one ASV classified as subcluster 5.1 (belonging to the better-studied marine-specific group) (Partensky et al. 1999; Callieri et al. 2013; Celepli et al. 2017; Xia et al. 2017) (**Fig. 5B**; **Table S1**: *Synechococcus* RA). We observed the *Synechococcus* subcluster 5.2 ASV3 at > 1% relative abundance at the majority of the sites sampled while we only found the ASV70 *Synechococcus* subcluster 5.1 at low relative abundances in salinities > 12. *Synechococcus* subcluster 5.2 ASV47 occurred in a limited number of samples (< 50%), but a broad range of salinities (**Fig. 5B**). ASV47 may be further selected for by season as relative abundances were > 0.1% during the late spring and summer months (**Fig. 5B**; **Table S1**: *Synechococcus* RA).

**Figure 5.**
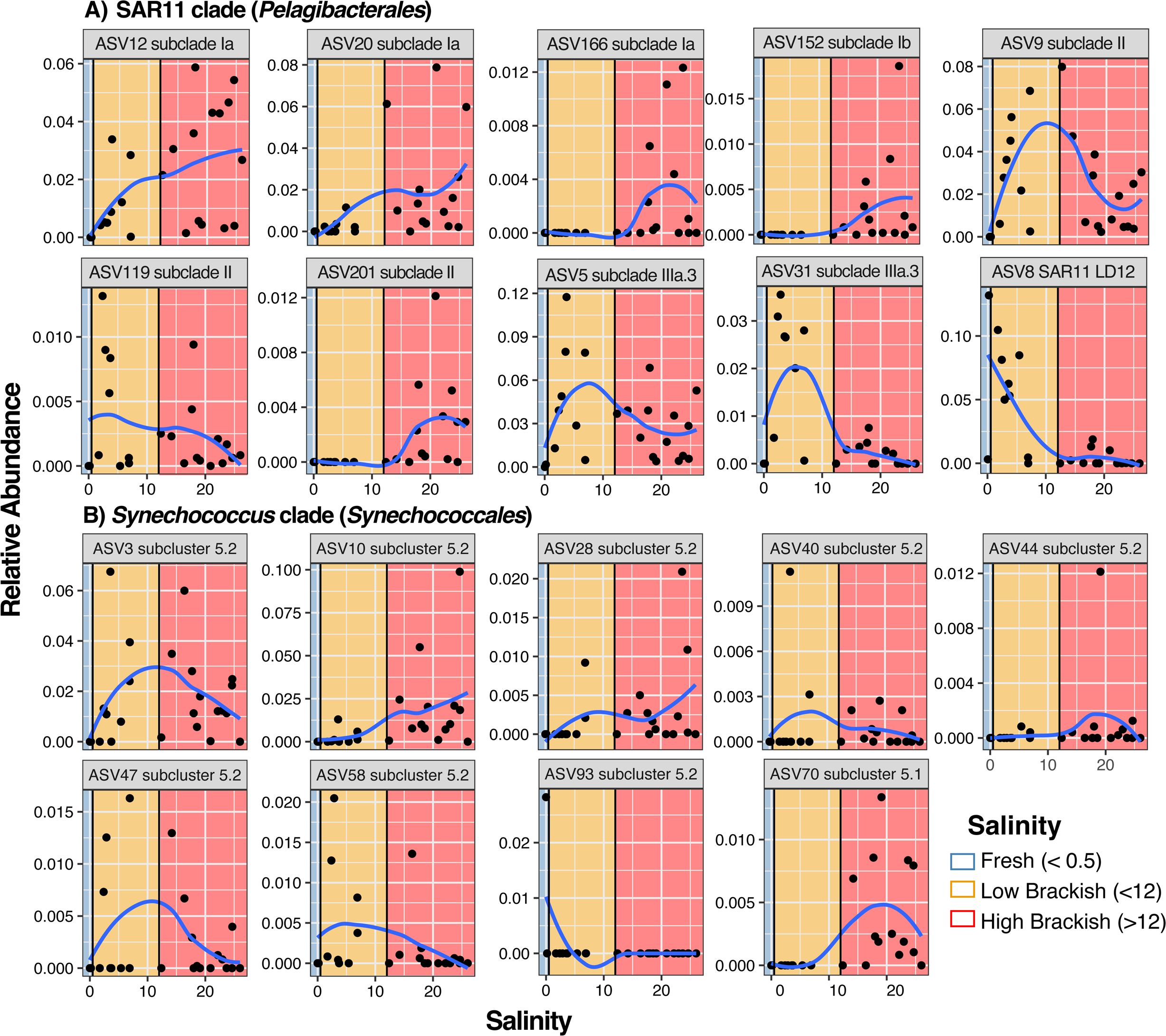
0.2-2.7 µm fraction ASV relative abundance within key taxonomic clades, A) SAR11 B) *Synechococcus*, across with salinity. Nonlinear regression lines are provided as a visual aid for abundance trends.

### > 2.7 µm associated communities

The > 2.7 µm size fraction communities were composed of bacteria classified as Cyanobacteria, Proteobacteria, Actinobacteria, and Bacteroidota (**Fig. 3B**, **Table S1**: > 2.7 µm RA). We observed numerous taxa known for planktonic lifestyles at high abundances in the > 2.7 µm size fraction (e.g, *Cyanobium* and the SAR11 clade; **Fig. S1**) (Partensky et al. 1999; Crump et al. 1999; Giovannoni 2017). The presence of typical planktonic taxa within the > 2.7 µm fraction complicates the interpretation of our results of the prefilter samples. While organisms may be present on this fraction due to cell size or their attachment to large particles, the presence of taxa such as the SAR11 and *Synechococcus* clades with known cell sizes of < 2.7 µm and planktonic lifestyles (Olson et al. 1990; Rappé et al. 2002; Garcia et al. 2016; Henson et al. 2018b; Lanclos et al. 2023) suggests that the high sediment loads found in the nGoM may have acted as an additional filter (**Figs. 3B, 4B**). Indeed, the amount of volume filtered can act as a secondary filter, trapping planktonic cells on the prefilter (> 2.7 µm fraction) and biasing downstream analyses (Padilla et al. 2015). While turbidity was not part of the environmental parameters we collected, high turbidity is common across the nGoM, owing to the influence of runoff and the Mississippi and Atchafalaya Rivers (Wiseman et al. 1999; Dodds 2006). Moreover, SAR11, Synechococcus, and SAR86-all known planktonic organisms-have been observed in particle-associated samples from the Amazon River plume (Satinsky et al. 2014b; a, 2017). Although we did not investigate the impact of filtered volume on which taxa are observed in the 2.7 µm filters, our results emphasize that researchers should consider sediment load, in addition to volume, when working in coastal and estuary environments with high turbidity.

The > 2.7 µm communities had a significantly higher species richness (**Fig. S2**) than the planktonic fraction, which is typical of particle-associated communities (Crump et al. 1999; D’Ambrosio et al. 2014). However, the incorporation of planktonic taxa into the larger filter fraction may have also increased the > 2.7 µm community richness. Statistically, the two size-fractionated communities were more strongly differentiated by salinity than filter fraction (Salinity: ANOSIM R=0.82, p=0.001, Filter: ANOSIM R=0.244, p=0.001), a result that may be partially explained by the presence of planktonic taxa in the > 2.7µm fraction. When we examined the > 2.7 µm communities alone, salinity and silicic acid were the most significant factors driving separation (Salinity: R^2^=0.812, p=0.001; Silicic acid: R^2^=0.686, p=0.001), with NO_2_ ^−^ (R^2^=0.482, p=0.006) and NH_4_ ^+^ (R^2^=0.354, p=0.022) also correlating to a lesser extent with the ordination (**Table S1**: envfit).

Despite our observation that numerous abundant ASVs in the > 2.7 µm fraction were associated with planktonic lifestyles, we also observed many typical sediment or particle-associated organisms throughout the top 25 rank abundance curve and across the majority of sites. These include ASV21 (unk. *Saprospiraceae*), ASV33 (*Pseudoalteromonas* sp.), ASV52 (unk. *Sphingobacteriales*), and ASV56 (unk. *Verrucomicrobiae*) (**Figs. 3B, S1**) (Mason et al. 2016; Milici et al. 2016; Mitulla et al. 2016; Satinsky et al. 2017; Liu et al. 2020). Like in the 0.2-2.7 µm fraction, many > 2.7 µm ASVs had differential abundances across salinity. We observed ASVs such as ASV17 (*Sediminibacterium* sp.), ASV77 and ASV114 (unk. *Kapabacteriales*), and ASV116 (unk. *Pedosphaeraceae*) enriched at freshwater sites, with ASV84 (unk. *Cryomorphaceae*) more abundant in high brackish waters (**Fig. 4B; Table S1**: Pairwise), supporting previous findings (Bowman 2011). Notably, the RS62 marine group of Betaproteobacteria (ASV25) was significantly enriched (max relative abundance=0.30) in the particle-associated fraction at site FWC in April 2016 (FWC2) (**Fig. 3B**). Outside of FWC2, it maintained a low relative abundance (median relative abundance=0, average relative abundance=0.01) (**Fig. 3B**; **Table S1**: >2.7 µm community), suggesting a potential bloom response. RS62 taxa occurred in high abundances in the planktonic fractions in the Pearl River estuary system (Liu et al. 2020), but had strong associations with phytoplankton blooms elsewhere (Francis et al. 2021), which may explain its abundance patterns in the > 2.7 µm fraction.

### Brackish communities

Although salinity was the strongest factor separating the microbial communities in our samples (**Fig. 2**; **Table S1**: envfit), many taxa occurred across all salinities with peak relative abundances at sites with brackish conditions (**Figs. 3, 5, S1**). This euryhaline distribution can signify a specifically brackish-water-adapted community. We observed many taxa with high relative abundance and frequency that were also poorly correlated to salinity (spearman rank correlation coefficient [rho] near 0) (**Fig. 6, S5**), supporting the conclusion of a brackish-adapted core microbiome in the nGoM estuaries. Taxa more negatively or positively correlated to salinity are more indicative of freshwater or marine-associated organisms, respectively. Within both filter fractions, the freshwater group contained abundant taxa from well-known groups such as *Candidatus* Fonsibacter sp. (SAR11 LD12) and the acI clade, whereas the high brackish group contained taxa that increased in abundance with salinity such as SAR11 subclade I, SAR324 clade, *Synechococcus* subcluster 5.2, and *Candidatus* Actinomarina, as well the MGII clade (*Thermoplasmatota*) in the > 2.7 µm fraction (**Fig. 6**). Some microorganisms in the brackish group maintained high median relative abundances and occurred at > 75% of the sites sampled over three years (**Fig. 6**), and therefore likely represent autochthonous brackish and/or euryhaline nGoM taxa. These abundant brackish taxa from both fractions included SAR11 subclade IIIa.3 (ASV5), OM43 (ASV7), Aegean-169 (ASV24), and SAR11 subclade II (ASV9) (**Fig. 6**, **Table S1**: Spearman Rank Correlations), as well as unk. *Saprospiraceae* (ASV645) and unk. Gammaproteobacterium PLTA13 (ASV95) in the > 2.7 µm fraction. We also note that SAR11 subclade IIIa.3 had an ASV that was more associated with lower salinity (ASV31-**Fig. 5A**), and thus salinity-based ecotypes may occur at the ASV level within some clades. Comparatively, the > 2.7 µm fraction had fewer autochthonous brackish ASVs, but a core group was still present (**Fig. 6B**, **Table S1:** Spearman Rank Correlations).

**Figure 6.**
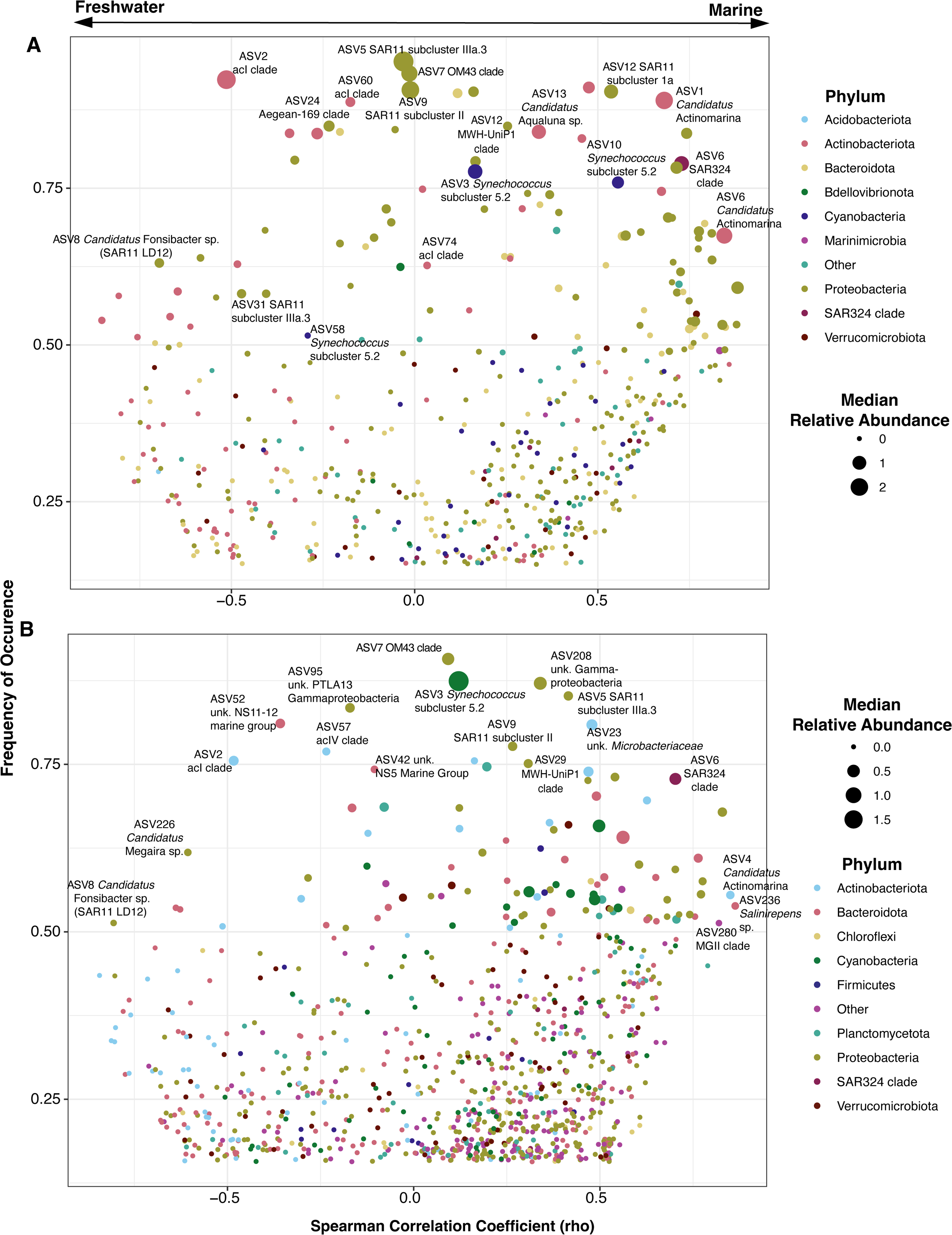
Two-sided Spearman’s rank correlation coefficient (rho) of 0.2-2.7 µm (A) and > 2.7 µm (B) fraction ASVs according to its frequency of occurrence across all sites. Only ASVs that appear in at least four sites are displayed. ASVs are color-coded by phylum and the size of the dot corresponds to the median relative abundance for each ASV.

## Discussion

Estuarine ecosystems have distinct and heterogeneous microbial community structures compared to open-ocean marine environments due to the contributions from both riverine and marine sources (Crump et al. 2004; Herlemann et al. 2014; Fortunato and Crump 2015; Hugerth et al. 2015; Wang et al. 2020). Although this observation has been reproduced in a number of locations, estuarine and coastal systems are drastically undersampled for microbial observation compared to open ocean environments, and the extent to which brackish microbiomes from these diverse estuary communities overlap is still unknown. This leaves a substantial knowledge gap that is critical for understanding global biogeochemical cycling because coastal communities can perform a disproportionate amount of turnover, for example with carbon cycling, compared to their open ocean counterparts (Cai 2011; Bauer et al. 2013). Furthermore, the interconnected coastal and estuarine ecosystems are under worsening pressure due to land loss and sea level rise (Nittrouer et al. 2012; Reed et al. 2020). The samples we collected across the nGoM Louisiana coastline represent a diverse collection of environments with notable differences in salinity and other environmental regimes compared to the more stable open ocean. Given the continual and projected worsening of land loss, oil spills, and natural disasters in this region, these data will serve as an important resource for future researchers investigating the microbial communities found across the coastal nGoM and coastal estuaries globally.

Salinity has repeatedly been observed as the strongest evolutionary and ecological selective factor across aquatic clades and ecosystems (Lozupone and Knight 2007; Logares et al. 2009, 2010; Dupont et al. 2014). The fluctuating salinity gradients within estuaries lead to transitory freshwater and marine lineages with fewer well-adapted brackish community members (Wang et al. 2020), generalized in the Remane curve relating richness to salinity (Remane 1934; Remane and Schlieper 1971; Olli et al. 2019). Although not originally formulated to describe microorganisms, the pattern applies to microbes, and the relationships described in the Remane curve anticipated the now supported observation of infrequent transitions between marine and freshwater species (Logares et al. 2009, 2010; Olli et al. 2019, 2022) and the importance of salinity in structuring microbial communities (Lozupone and Knight 2007; Logares et al. 2009; Walsh et al. 2013; Dupont et al. 2014; Paver et al. 2018). Indeed, typical freshwater taxa such as acI and *Candidatus* Fonsibacter sp. (SAR11 LD12) taxa were strongly correlated with sites with salinities below five while typical marine taxa were more strongly correlated with sites with salinities above twenty, highlighting the freshwater marine divide and the dynamic nature of coastal estuaries found in other studies (Campbell et al. 2011; Campbell and Kirchman 2012; Ghylin et al. 2014; Herlemann et al. 2014; Henson et al. 2018b; Rasmussen et al. 2020; López-Pérez Mario et al. 2020).

However, sites such as Bay of Batiste (BBAT) and Freshwater City (FWC) have large variations in salinity (oligohaline to polyhaline), which had a large impact on community composition (**Fig. 2**) such as switching taxa from Actinobacteria acI to OM1 or SAR11 subclade IIIa to subclade I. Shifts in these distinct communities would have important cellular energetics implications such as change in central metabolism: Embden-Meyerhof-Parnas glycolysis (freshwater) versus Entner–Doudoroff pathway (marine), loss of C1 metabolism (freshwater), and the reliance on de novo synthesis (freshwater) rather than uptake (marine) of many important amino acids, osmolytes and other compounds (Dupont et al. 2014; Henson et al. 2018b; Lanclos et al. 2023; Jurdzinski et al. 2023). Changes in these energetics and metabolisms, de novo synthesis versus uptake or loss of C1 metabolism could fundamentally alter carbon cycling and other nutrient availability. Therefore, considering the variability in energetics alongside nutrient availability over time and space is critical for effective future conservation and management strategies to consider under future climate and sea-level rise scenarios.

Taxa like SAR11 subclade IIIa, OM43 clade, and *Synechococcus* subcluster 5.2 have been previously established as core members of the brackish-water microbiome (Chauhan et al. 2009; Larsson et al. 2014; Herlemann et al. 2014; Hugerth et al. 2015; Vidanage et al. 2020; Cabello-Yeves et al. 2022; Lanclos et al. 2023; Jurdzinski et al. 2023). Our community analysis fortifies these assignments and also highlights the inclusion of other taxa such as SAR11 subclade II, the acI C2 tribe, SAR324, the MWH-UniP1 clade (ASV29), and specific ASVis within poorly classified groups like unk. *Flavobacteriales* spp. (ASV42, ASV94) and unk. *Planctomycetota* (ASV92) among core brackish-water members (**Fig. 2, 4, 6**). Many of these groups are typically considered “marine” or “freshwater” taxa. Nevertheless, in addition to the nGoM, these taxa have been observed in brackish environments such as the Baltic Sea, Chilika Lagoon (Bay of Bengal), and the Chesapeake and San Francisco Bays (Campbell and Kirchman 2012; D’Ambrosio et al. 2014; Hugerth et al. 2015; Mohapatra et al. 2020; Rasmussen et al. 2020; Wang et al. 2020; Getz et al. 2023). Thus, our study expands the known biogeography and salinity ranges of these organisms, and provides additional evidence of taxa within these clades with potential preference for brackish environments, not just low salinity or marine. SAR11 subclade II is found predominantly in temperate open oceans during deep mixing events (Vergin et al. 2013) and in oxygen minimum zones (Tsementzi et al. 2016). However, this subclade was also recently established as highly abundant in brackish environments (Lanclos et al. 2023). Phylogenetically, the two abundant brackish subclade II ASVs ASV9 and ASV119 formed a unique cluster of nGoM taxa and were sister to taxa found in the San Francisco and Chesapeake Bays (**Fig. S3**). Our results support the hypothesis that a subset of the SAR11 subclade II is brackish-adapted with potentially unique distributions, similar to other ecotypes within SAR11 such as subclade IIIa (Campbell et al. 2022; Lanclos et al. 2023). While salinity transitions are thought to be rare within a group (Logares et al. 2009, 2010), the distribution and population size of the SAR11 clade may have facilitated multiple transitions to brackish intermediaries within the different subclades. Moreover, whether the nGoM cluster of ASVs represents a unique tribe of brackish-adapted ecotypes in subclade II or is part of a large cluster of brackish organisms is still unknown, but highlights the missing diversity within an important aquatic group (Campbell et al. 2022; Lanclos et al. 2023) and the need for additional in-depth sampling of estuary ecosystems.

While the majority of ASVs from the acI Actinobacteria were strongly correlated with low salinity and freshwater environments, two ASVs (ASV60, ASV74) were uncorrelated with salinity. Unlike other acI freshwater ASVs, ASV60 and ASV74 both had maximum abundance at intermediate salinities, suggesting these taxa may represent brackish-adapted subclade within the acI clade. Phylogenetically, ASV74 clusters with other nGoM taxa sister to taxa within the acI-C2 tribe while ASV60 clusters with other nGoM ASVs as an early diverging member within the acI clade (**Fig. S4**). Previous studies have observed taxa within the “freshwater” acI at salinities of up to 14 (Holmfeldt et al. 2009; Hugerth et al. 2015; Mehrshad et al. 2016). We provide strong evidence of taxa within acI-C with unique distributions at brackish salinities across multiple GoM estuaries, suggesting members of the acI clade may be core members of both fresh and brackish environments (**Table S1**: 0.2-2.7 µm RA). However, given our limited taxonomic and metabolic clarity with amplicon sequencing, further genomic and physiological studies are needed to place these taxa within the acI clade and test their salinity preferences. Taken together with the distribution of other microbial clades, our data suggest that brackish environments are important diversification hotspots for aquatic microorganisms, which may result from the continuous exposure to fluctuating salinity gradients.

The SAR324 clade (previously known as Marine Group B, as a member of the Deltaproteobacteria, and now as a candidate phylum) is found throughout the global oceans but predominantly in bathypelagic waters (Haroon et al. 2016; Boeuf et al. 2021; Malfertheiner et al. 2022). Nevertheless, the SAR324 clade has also been observed as a significant member of planktonic estuarine and coastal communities, for example in the Amazon River plume, as well as in other estuaries and coastal sediment (Satinsky et al. 2017; Boeuf et al. 2021; Malfertheiner et al. 2022). Within our coastal nGoM samples, the SAR324 clade was the 12th most abundant taxon (ASV6) and was present in all samples with a salinity > 2 (**Figs. 3A, S1**), peaking in relative abundance between salinities of 18-23 before sharply decreasing (**Table S1**: 0.2-2.7 µm RA**, Fig. S1**). ASV6 and others (e.g., ASV328, ASV653, ASV690) classified as SAR324 imply a brackish-adapted subclade within SAR324 that warrants further investigation (**Table S1**: 0.2-2.7 µm RA). Organisms within the SAR324 clade possess the potential for flexible metabolic capacity such as the ability to utilize complex organics, light, as well as fix carbon via the Carbon-Benson-Basson cycle (Malfertheiner et al. 2022; Jurdzinski et al. 2023), which may facilitate their ecological and evolutionary transition to brackish environments. Previous phylogenomic studies on cross-biome transitions found that plasticity may be a hallmark feature of brackish-adapted organisms (Jurdzinski et al. 2023). Future studies of this clade should incorporate more estuarine samples to help further delineate if it is part of the global brackish microbiome and investigate the functional diversity and processes that led to its differentiation.

Without physiological testing, it is difficult to ascertain the specific salinity preference of a taxon, e.g. fresh, brackish, or marine. An organism may be euryhaline, capable of growth across a wide range of salinities, but not brackish-adapted, despite both euryhaline and brackish-adapted organisms sharing adaptive salinity responses. For instance, while cultivars from SAR11 subclade IIIa.3 and IIIa.1 are both euryhaline, the IIIa.3 cultivar had optimal growth in mesohaline salinities, while IIIa.1 grew similarly across salinities >10 (Lanclos et al. 2023). Similarly, two coastal isolates within the euryhaline *Synechococcus* subcluster 5.2 CB4 clade had varying growth optimums driven by differences in their capacity to regulate osmolyte production and metabolic capacity (Xia et al. 2017). Therefore, it is important for future studies to examine the physiological characteristics of taxa within the pan-brackish microbiome to unveil the genomic underpinnings that delineate euryhaline and brackish-adapted taxa.

A major and growing concern for many stakeholders in freshwater and coastal environments is harmful algal blooms (HABs). Analagous to red tide HABs in marine environments, cyanobacterial HABs (cyanoHABs) occur in freshwater and can lead to fish kills, contaminated drinking water, and economic and ecological loss (Carmichael 2001; Backer 2002; Anderson 2009; Roy et al. 2013; Chaffin et al. 2021). Within the > 2.7 fraction, two taxa, ASV77 and ASV114 (unk. *Kapabacteriales),* were significantly correlated with low salinity (**Fig. 4B**; **Table S1**: Pairwise), the 2^nd^ and 4^th^ most abundant taxa at Lake Martin, and blasted to numerous OTUs (BLASTn hits > 99%) from eutrophic freshwater lakes and HABs (Cai et al. 2013), particularly *Microcystis* blooms (Li et al. 2012). Furthermore, a recently assembled metagenome-assembled genome from the *Kapabacteriales* originated from a culture of the HAB-causing cyanobacterium *Dolichospermum* (Al-Saud et al. 2020). Thus, at least some Kapabacteriales are closely associated with cyanoHABs and may be indicators of such. Within both fractions, numerous other ASVs associated with cyanoHABs (e.g., *Microcystis* and *Planktothrix*) were found at freshwater coastal (ARD) and inland (Lake Martin) sites, albeit at lower abundances than the *Kapabacteriales*-associated ASVs. Although no cyanoHABs were reported during the time of sampling at these sites, the presence of these cyanobacteria and other associated bacteria highlights the potential for cyanoHABs to impact these coastal locations, particularly at lower salinity sites like ARD (Roy et al. 2013).

### Conclusion

Estuarine and coastal ecosystems are diverse environments influenced by tidal fluxes, interconnected wetlands, and river outflows with high economic and ecological importance (Rabalais et al. 1996; Adams et al. 2004; Barbier et al. 2011; Thrash et al. 2017). Our study of the microbial communities from the nGoM Louisiana coastline makes an important contribution to the aquatic microbial ecology of these understudied habitats that represent a diverse collection of environments with notable differences in salinity and other environmental characteristics. Our data contributes to the growing knowledge of the globally distributed, core brackish microbiome, composed of members from important aquatic clades such as OM43, *Synechococcucs*, and SAR11 (Larsson et al. 2014; Hugerth et al. 2015; Celepli et al. 2017; Vidanage et al. 2020; Wang et al. 2020; Cabello-Yeves et al. 2022; Lanclos et al. 2023; Jurdzinski et al. 2023) that should be expanded to include taxa from groups such as acI Actinobacteria, MWH-UniP1 Betaproteobacteria, SAR324, and SAR11 subclade II. Moreover, this study highlights the potential for estuaries to house important biodiversity, such as the observed brackish cluster within the freshwater acI clade. Future research should incorporate time series data, collection of genomic information, and new culturing efforts to help resolve population-level diversity, function, and temporal variation of the endogenous brackish community in these dynamic environments.

## Methods and Materials

### Sample Collection

Surface water (< 1 m) was collected at nine different sites. Sampling was done as previously described (Henson et al. 2016, 2018b, 2020). Briefly, duplicate 120 mL water samples were sequentially filtered through a 2.7 µm GF/D filter (Whatman, UK) and 0.2 µm Sterivex filter (Millipore, USA) using a handheld 60 mL syringe (Becton-Dickinson, USA. Sterivex filtrate was analyzed for SiOH_4_, NO_3_ ^−2^, NO_2_ ^−^, NH_4_ ^+^, and PO_4_ ^3-^ at the University of Washington Marine Chemistry Laboratory (http://www.ocean.washington.edu/story/Marine+Chemistry+Laboratory). Filters were immediately placed on ice, transferred to the lab (maximum of 3 h on ice), and frozen at −20℃ until further processing. Baseline water conditions of temperature, salinity, pH, and dissolved oxygen were measured using a handheld YSI 556 multiprobe system (YSI Inc., USA). All site locations (latitude and longitude), water chemistry, water conditions, and sampling dates can be found in **Table S1**.

### nGoM coastal sites

Sites were sampled once a year for three years, except for Terrebonne Bay, which was collected twice. The sites sampled were Lake Borgne (LKB, Shell Beach, LA), Bay Pomme d’Or (JLB, Buras, LA), Terrebonne Bay (TBON, Cocodrie, LA), Atchafalaya River Delta (ARD, Franklin, LA), Freshwater City (FWC, Kaplan, LA), and Calcasieu Jetties (CJ, Cameron, LA) (**Table 1**). The sample collection was done previously as part of the three-year cultivation campaign between September 2014 and February 2017 (Henson et al. 2016, 2020).

### Estuarine nGoM sites

Two sites were sampled once roughly every two months for five months between July, 2015 and November 2015. The sites sampled were Sabine Wetlands (Sabine, Cameron, LA) and Bay Batiste (BBAT, Port Sulphur, LA) (**Table 1**). Water was collected from the surface water (< 1 m) and filtered immediately on site as described above.

### Lake Martin

Lake Martin (Swamp, Beaux Bridge, LA) was sampled as an inland lake representative in November 2014 (**Table 1**). Water was collected from the surface water (< 1 m) and filtered immediately on site as described above.

### DNA Extraction and Sequencing

In total, 96 samples were collected, extracted for DNA, and sequenced. All DNA was extracted and sequenced as previously described (Henson et al. 2020). Briefly, DNA from both size fractions were extracted using the MoBIO PowerWater DNA kit (MoBIO, USA) and quantified using the Qubit high-sensitivity dsDNA assay kit (ThermoFisher, USA). DNA was frozen at −20°C until sequencing. DNA was sequenced targeting the 16S rRNA gene V4 region with the 515F-806R primer set (Apprill et al. 2015; Walters et al. 2016) at Argonne National Laboratory using Illumina MiSeq 2 x 250 bp paired-end reads.

### Data Analyses

In total, 9,655,001 raw reads were obtained from sequencing. Data was curated and processed in R (v4.2.2) with the package *DADA2* (v1.21.0) following their published protocol (Callahan et al. 2016). Briefly, data were preprocessed for quality and length using *filterAndTrim* with the flags truncLen=c(240,160), maxN=0, maxEE=c(2,2), truncQ=2, rm.phix=TRUE, compress=TRUE. Using the error rates calculated from the *learnErrors,* unique sequences were determined and the resulting reads were merged using *mergePairs*. Sequences shorter than 250 and larger than 256 were removed. Chimeras were removed using the tool *removeBimeraDenovopoor* with the method flag “consensus”. Taxonomy was assigned using the Silva V132 training set and the *assignTaxonomy* command (Pruesse et al. 2007; Quast et al. 2013).

Because of poor sequencing depth (< 1000 sequences), two samples from the > 2.7 µm fraction from the Bay Batiste (BBAT1A PRE and BBAT1B PRE) were removed from all downstream analyses, resulting in 94 remaining samples. ASVs assigned to Chloroplast (Order), Mitochondria (Family), Eurkaryote (Kingdom), and unknown (Kingdom) were removed manually. Data were processed using the program PhyloSeq in the R (v4.0.2) statistical environment following a protocol similar to those previously published (Henson et al. 2018a; b, 2020). Our modified PhyloSeq script is available on our GitHub repository, https://github.com/thrash-lab/Modified-Phyloseq. After filtering, the data set contained 11,791 unique ASVs and 7,341 ASVs after rarefying with the function *rarefy_even_depth* (**Table S1**). Alpha diversity was calculated on unrarefied data using the function *plot_richness*. Statistical comparisons between size fraction alpha diversity were made using the function *geomsignif* from the package ggsignif (Ahlmann-Eltze and Patil 2021). Beta diversity between sites was examined using Bray-Curtis distances via ordination with nonmetric multidimensional scaling (NMDS). Measured environmental parameters were normalized using the R function *scale* before downstream analyses. The significance of the transformed environmental parameters on beta diversity was calculated with the *envfit* function (Oksanen et al. 2015). Relative abundances of an ASV from each sample were calculated and averaged between biological duplicates as previously published (Henson et al. 2018a, 2020).

### acI and SAR11 taxonomic classification

To better resolve the taxonomic and ecotype designation of acI and SAR11 clade-classified ASVs, ASVs were clustered with near full-length 16S rRNA gene sequences. Sequences were obtained from Lanclos et al. 2023 for SAR11 and from Holmfeldt et al. 2009 and TaxAss for acI (Rohwer et al. 2018). ASV phylogenetic placement was inferred as previously described (Henson et al. 2020; Lanclos et al. 2023). ASVs with improved taxonomic resolutions were updated with new clade designations (**Fig. S3 and S4**).

### Statistical Analyses

All statistical analyses were performed in R (v4.2.2). Water chemistry for principle components analysis was normalized using root-mean-square with the command *scale()* and then conducted using the *rda()* function. For comparing non-normally distributed data (relative abundance versus salinity group), a Kruskal Wallis test followed by a Mann-Whitney U test was performed (Schloss Patrick D. 2023). We employed a two-sided Spearman’s rank correlation to determine the relationship between taxa relative abundance and salinity. To control for the false discovery rate, p values were adjusted using the Benjamini & Hochberg method (Benjamini and Hochberg 1995) using *p.adjust()* or as a flag. Our modified scripts are available on the GitHub repository, https://github.com/theaquaticmicrobiologylab/Henson_Thrash_CoastalnGoM.

## Data Records

The *nGoM coastal sites* 0.2-2.7 µm fraction iTag sequences are available at the Sequence Read Archive (SRA) with accession numbers SRS2644905 - SRS2644921 as previously published (Henson et al. 2018b, 2020). All other 0.2-2.7 µm fraction iTag sequences are available at the SRA with accession numbers SRS1840441 - SRS1840447. All > 2.7 µm fraction iTag sequences are available at the SRA with accession numbers SRR18184264 - SRR18184311.

## Acknowledgments

A portion of this research was conducted with high-performance computing resources provided by Louisiana State University (http://www.hpc.lsu.edu) and the authors acknowledge the Center for Advanced Research Computing (CARC) at the University of Southern California for providing computing resources that have contributed to the research results reported within this publication. URL: https://carc.usc.edu.This work was funded by the Louisiana Board of Regents (Board of Regents) (LEQSF[2014-2017]-RDA-06), the Louisiana State University Department of Biological Sciences, a National Academies of Science, Engineering, and Medicine Gulf Research Program Early Career Research Fellowship, and a Simons Foundation Early Career Investigator in Marine Microbial Ecology and Evolution Award to J.C.T.; the Dornsife College of Letters, Arts, and Sciences at the University of Southern California; the Department of Biological Sciences and College of College of Liberal Arts and Sciences; and a LEEC grant from the Louisiana Department of Fish and Wildlife, Lerner Gray grant from the American Natural History Museum, and a Simons Foundation Marine Microbiology Postdoctoral Fellowship to M.W.H..

## Supplemental Figure

**Figure S1.** Rank abundances of the 25 most abundant ASVS from all sites based on the average relative abundance in the 0.2 - 2.7 µm fraction (A) and > 2.7 µm fraction (B). The boxes indicate the interquartile range (IQR) of the data, with vertical lines indicating the upper and lower extremes according to 1.5 × IQR. Horizontal lines within each box indicate the median. The data points comprising the distribution are plotted on top of the boxplots. The color of the dot represents the broad salinity classification of fresh (blue, < 0.5 salinity), low salinity (orange, < 12 salinity), and high salinity (red, > 12 salinity). The black dot is the average relative abundance across all sites.

**Figure S2.** Alpha diversity was calculated for the 0.2 - 2.7 µm and > 2.7 µm size fractions. The boxes indicate the interquartile range (IQR) of the data, with vertical lines indicating the upper and lower extremes according to 1.5 × IQR. Horizontal lines within each box indicate the median. The data points comprising the distribution are plotted on top of the boxplots. Asterisks indicate the strength of significance as calculated using a one-way ANOVA (“***”=0.001,“**”=0.01,“*”=0.05).

**Figure S3.** 16S rRNA gene phylogeny of the SAR11 clade with nGoM ASVs. The scale bar represents 0.1 changes per position. Values at internal nodes indicate bootstrap values (n = 1000).

**Figure S4.** 16S rRNA gene phylogeny of the acI clade with nGoM ASVs. The scale bar represents 0.1 changes per position. Values at internal nodes indicate bootstrap values (n = 1000).

**Figure S5.** The relative frequency of ASVs by the observed two-sided Spearman’s rank correlation coefficient (rho) from the 0.2-2.7 µm (A) and > 2.7 µm (B) fraction communities. Two-sided rho values were rounded to the nearest quarter decimal. Between the red dashed lines are taxa poorly correlated (−0.25 to 0.25) to salinity. A blue nonlinear regression line is provided as a visual aid for trends.

## Supplemental Table

Table S1. A spreadsheet with multiple tabs including supplemental data and results. Table S1 is hosted through FigShare, https://figshare.com/projects/Microbial_ecology_of_coastal_northern_Gulf_of_Mexico_waters/186288.

