## Supplementary figures and images for "Microbial ecology of coastal northern Gulf of Mexico waters"

### Supplemental Figure 1

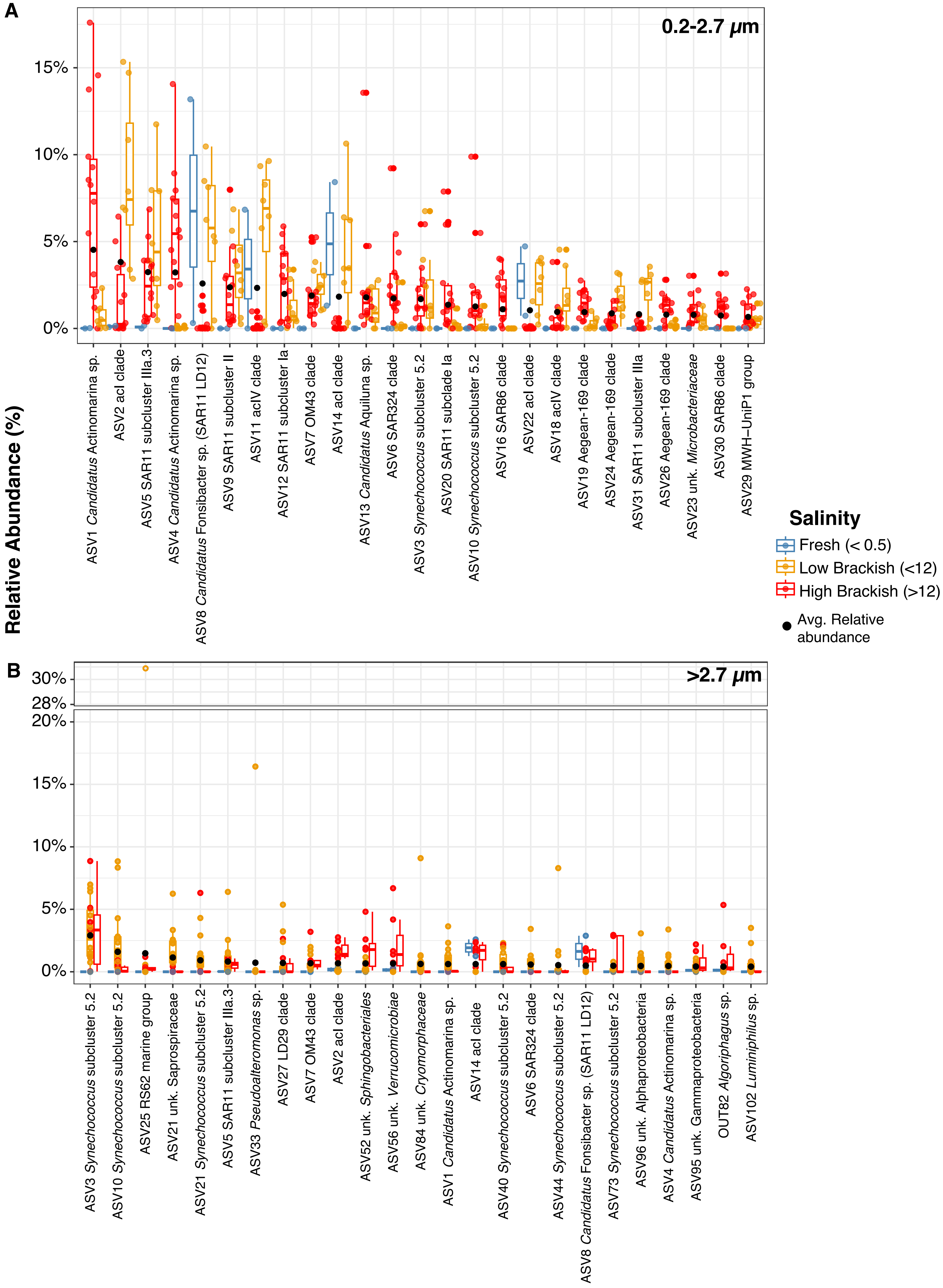

### Supplemental Figure 2

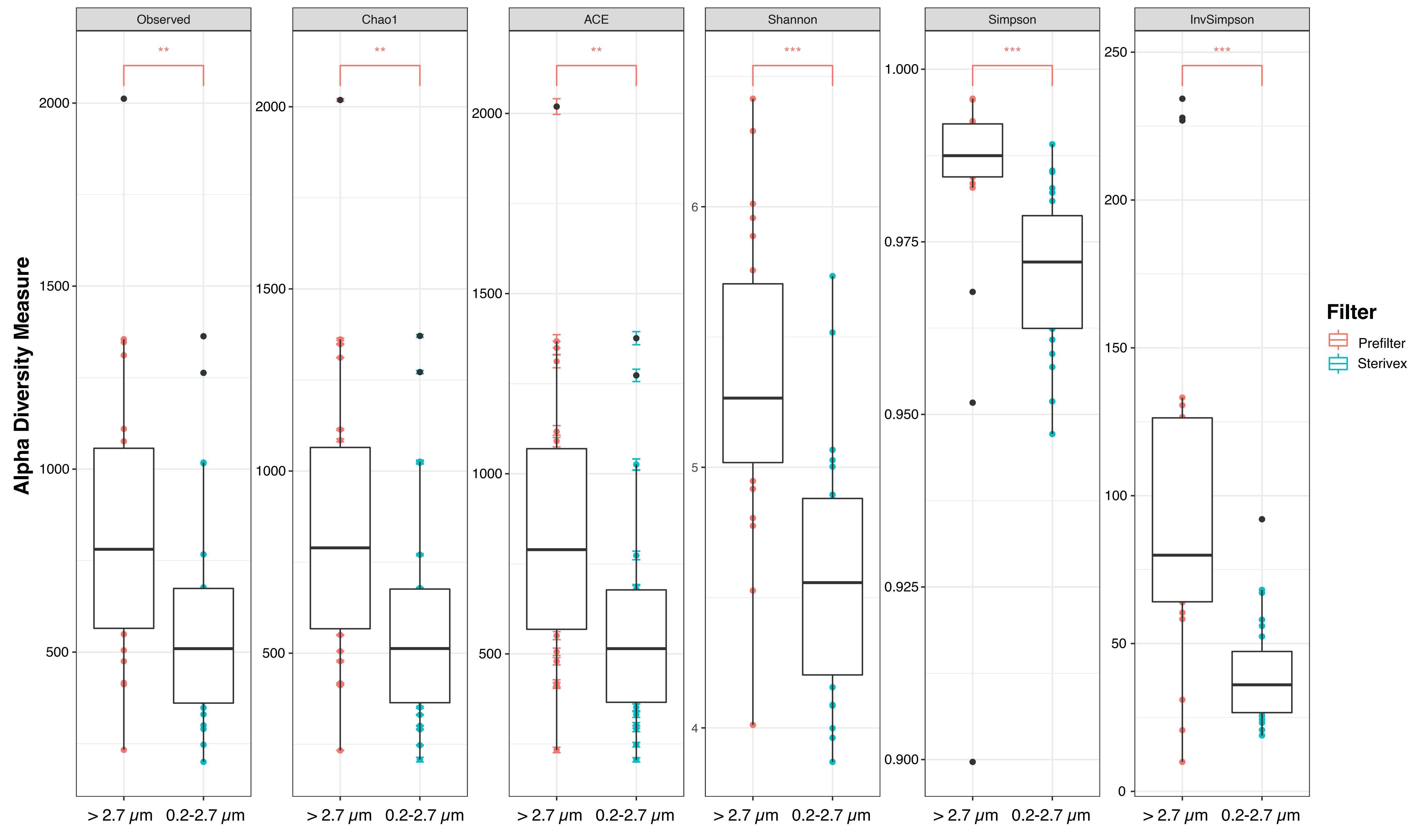

### Supplemental Figure 3

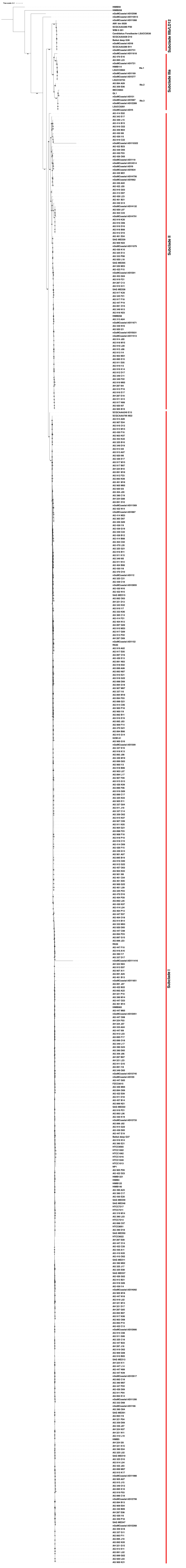

### Supplemental Figure 4

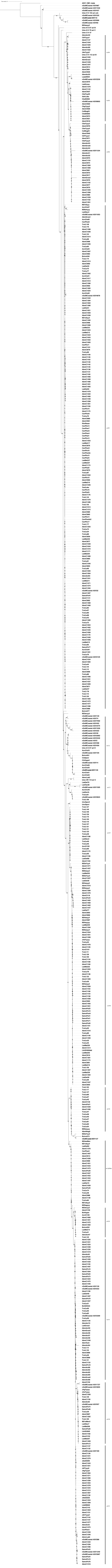

### Supplemental Figure 5

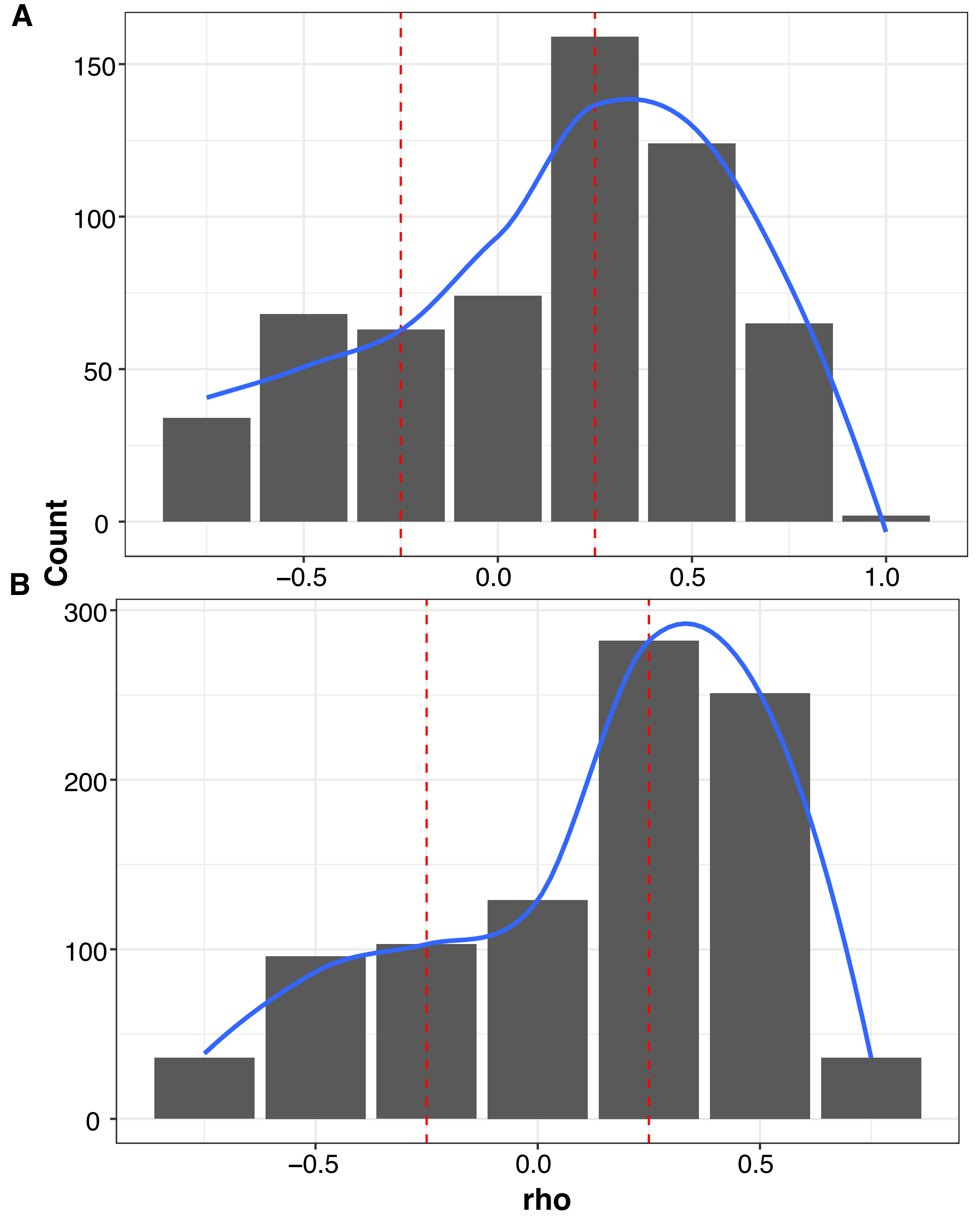
